# Computational neuroanatomy of human stratum proprium of interparietal sulcus

**DOI:** 10.1101/126961

**Authors:** Maiko Uesaki, Hiromasa Takemura, Hiroshi Ashida

## Abstract

Recent advances in diffusion-weighted MRI (dMRI) and tractography have enabled identification of several long-range white matter tracts in the human brain. Yet, our understanding of shorter tracts, such as those within the parietal lobe, remains limited. Over a century ago, a tract connecting the superior and inferior parts of the parietal cortex was identified in a post-mortem study: *Stratum proprium of interparietal sulcus* (SPIS; Sachs, 1892). The tract has since been replicated in another fibre dissection study (Vergani et al. 2014), however, it has never been identified in the living human brain and its anatomical properties are yet to be described. We used dMRI and tractography to identify and characterise SPIS in vivo, and explored its spatial proximity to the cortical areas associated with optic-flow processing using fMRI. SPIS was identified bilaterally in all subjects, and its anatomical position and trajectory are consistent with previous post-mortem studies. Subsequent evaluation of the tractography results using linear fascicle evaluation and virtual lesion analysis yielded strong statistical evidence for SPIS. We also found that SPIS endpoints are adjacent to the optic-flow selective areas. In sum, we show that SPIS is a short-range tract connecting the superior and inferior parts of the parietal cortex, wrapping around the intraparietal sulcus, and that it may be a crucial anatomy underlying optic-flow processing. In vivo identification and characterisation of SPIS will facilitate further research on SPIS in relation to cortical functions, their development, and diseases that affect them.

## 1. Introduction

Anatomical connections through the white matter axon bundles (i.e. fascicles, tracts) establish fundamental features of the brain’s information processing (Catani and Ffytche 2005; Catani and Thiebaut de Schotten 2012; Bullock et al. 2005; Fields 2008a, 2008b, 2015; Wandell and Yeatman 2013; Wandell, 2005). Diffusion-weighted magnetic resonance imaging (dMRI) and tractography provide a unique opportunity to identify and characterise the white matter tracts in the living human brain (Catani et al. 2002; Wakana et al. 2004; Mori and Zhang 2006; Catani and Thiebaut de Schotten 2012; Craddock et al. 2013; Wandell 2016).

A body of dMRI research has successfully identified several major long-range white matter tracts, such as the superior longitudinal fasciculus and the inferior longitudinal fasciculus, in a consistent manner with the known post-mortem anatomy (Catani et al. 2002; Wakana et al. 2004; Schmahmann et al. 2007), and ergo opened new avenues to research on the property of major human white matter tracts in relation to development and diseases (Lebel et al. 2012; Yeatman, Dougherty, Ben-Shachar, et al. 2012; Ogawa et al. 2014; Yeatman, Wandell, et al. 2014). More recent dMRI studies have identified shorter white matter tracts including frontal aslant tract (FAT) and vertical occipital fasciculus (VOF), which, partially for their relatively short trajectories, had previously received very little attention in the neuroscience literature (Catani et al. 2012; Catani et al. 2013; Yeatman et al. 2013; Yeatman, Weiner et al. 2014; Takemura, Rokem et al. 2016). Some of those studies have suggested the importance of the shorter tracts in relation to cognitive functions (Kinoshita et al. 2015; Kemerdere et al. 2016; Kronfeld-Duenias et al. 2016; Duan et al. 2015; Takemura, Rokem et al. 2016; Lee Masson et al. 2016).

Here, we focus on a short white matter tract connecting the superior and inferior parts of the parietal cortex, wrapping around the inferior parietal sulcus. This tract was initially described by the German neurologist Heinrich Sachs (1892) as *the stratum proprium of interparietal sulcus* (hereafter, we refer to this tract as SPIS). Except for one recent fibre dissection study replicating Sachs’s work (Vergani et al., 2014), this tract has been largely overlooked in the literature. Given the functional MRI (fMRI) evidence indicating the involvement of the superior and inferior parts of the parietal cortex in crucial cognitive functions (Corbetta and Shulman 2002; Uncapher and Wagner 2009; Cardin and Smith 2010; Blanke 2012; Greenlee et al. 2016), SPIS is a necessary and important tract supporting those functions. Yet, the characteristics of SPIS are poorly understood due to the lack of studies replicating SPIS in the living human brain, using three-dimensional digital anatomical data such as dMRI and reproducible computational analyses.

One of the cortical functions that involve the parietal cortex is optic-flow processing. Optic flow is the pattern of visual motion signals elicited by self-motion (Gibson 1950, 1954), and is an important cue to accurate perception of self-motion. A network of sensory areas in the parietal cortex has been shown to be involved in optic-flow processing (Cardin & Smith 2010, 2011). Those optic-flow selective areas include the ventral intraparietal area (VIP), the precuneus motion area (PcM) and the putative area 2v (p2v) located in the superior part of the parietal cortex, and the posterior-insular complex (PIC+: Deutschländer et al. 2004; Cardin and Smith 2010, 2011; Biagi et al. 2015; Uesaki and Ashida 2015; Wada et al. 2016) in the inferior part of the parietal cortex. Several studies have described a convergence of visual and vestibular information regarding self-motion, involving those optic-flow selective areas (Kleinschmidt et al. 2002; Wiest et al. 2004; Kovács et al. 2008; Fetsch et al. 2009; Butler et al. 2010; Prsa et al. 2015; Uesaki and Ashida 2015). In order to fully understand the underlying mechanism of optic-flow processing, it is essential to investigate how communication between the superior and inferior parts of the parietal cortex is supported by the white matter anatomy.

We used dMRI and tractography to identify the anatomical location and trajectory of SPIS in the living human brain. Ensemble tractography (Takemura, Caiafa, et al. 2016) yielded bilateral identification of SPIS in all 10 subjects. Evidence for SPIS was evaluated based on the consistency across datasets, comparison with post-mortem fibre dissection studies (Sachs 1892; Vergani et al. 2014), and virtual lesion analysis (Pestilli et al. 2014; Leong et al. 2016; Gomez et al. 2015; Takemura, Rokem et al. 2016). We also explored the functional relevance of SPIS by performing fMRI experiments on the same subjects and examining the spatial proximity between the SPIS endpoints and functionally-defined cortical regions previously associated with optic-flow processing (Cardin & Smith, 2010, 2011; Greenlee et al. 2016). Results showed that the dorso-lateral SPIS endpoints are near VIP, PcM and p2v, whereas the ventro-medial SPIS endpoints are near PIC+; placing SPIS in a good position to channel sensory signals between the distant cortical areas underlying visuo-vestibular integration necessary for optic-flow processing and perception of self-motion.

For the first time in the living human brain, our findings show that SPIS is a short-range tract connecting the superior and inferior parts of the parietal cortex, wrapping around the intraparietal sulcus. The findings also highlight that *in vivo* identification and characterisation of SPIS open new avenues to studying this tract in relation to cortical functions, their development, and diseases that affect them.

To facilitate future research, we have made the code to identify human SPIS publicly available [URL will be inserted upon acceptance].

## 2. Materials and methods

We analysed dMRI data of 10 human subjects, from two independent datasets. One set of dMRI data (KU dataset) was acquired at Kokoro Research Center, Kyoto University along with fMRI measurements to identify cortical regions activated by optic-flow stimulation. The other dMRI dataset was obtained from the Human Connectome Project (HCP dataset; Van Essen et al. 2013).

### 2.1. Subjects: KU Dataset

Six healthy volunteers (three males and three females; of the ages between 22 and 47 years; subjects S1-S6) participated in the study. All six subjects underwent both dMRI and fMRI experiments. All had normal or corrected-to-normal vision. All individual subjects gave written informed consent to take part in this study, which was conducted in accordance with the ethical standards stated in the Declaration of Helsinki and approved by the local ethics and safety committees at Kyoto University.

### 2.2. Data acquisition and preprocessing: KU Dataset

All MR images were obtained with a 3-Tesla Siemens Magnetom Verio scanner (Siemens, Erlangen, Germany), using a Siemens 32-channel head coil, at Kokoro Research Center, Kyoto University.

### 2.2.1. Diffusion-weighted MRI data

Two repeated acquisitions of MR images were conducted for each subject, using a diffusion-weighted single-shot spin echo, echo-planar sequence (60 axial slices, 2-mm isotropic voxels, time of repetition (TR) = 8300 ms, time echo (TE) = 94 ms, field of view (FoV) = 200 x 200 mm^2^). The dMRI data were sampled in 64 directions with a b-value of 1000 s/mm^2^. Two non-diffusion weighted images (b=0) were obtained.

Diffusion data were preprocessed using mrDiffusion, implemented in Matlab as part of the mrVista software distribution (https://github.com/vistalab/vistasoft). MR images in each scan were motion-corrected to the b = 0 image acquired in the same scan, using a rigid-body alignment algorithm implemented in SPM (Friston and Ashburner 2004). Eddy-current correction and head-motion correction were applied in the process of 14-parameter constrained nonlinear coregistration, based on the expected patterns of eddy-current distortion given the phase-encoding directions of the acquired data (Rohde et al. 2004). The gradient direction in each diffusion-weighted volume was corrected using the rotation parameters from the motion correction procedure. Subsequently, fibre orientation in each voxel was estimated using constrained spherical deconvolution (CSD; Tournier et al. 2007; L_max_ = 8) implemented in mrTrix (Tournier et al. 2012). CSD allows for tractography based on a model that is capable of accounting for crossing fibres. Because the tract connecting the superior and inferior parts of the parietal cortex likely intersects with the neighbouring major fasciculi such as the arcuate fasciculus, CSD was employed to fully reconstruct the tract.

### 2.2.2. T1-weighted image data

For each subject, a high-resolution T1-weighted 3D anatomical image was acquired with a magnetisation-prepared rapid-acquisition gradient echo (MP-RAGE; 208 axial slices, 1-mm isotropic voxels, TR = 2250 ms, TE = 3.51 ms, FoV = 256 x 256 mm^2^, flip angle = 9°, bandwidth = 230 Hz/pixel). Subsequently, the border between the grey and the white matters was defined. Initial segmentation was performed using an automated procedure in Freesurfer (Fischl 2012), which was then refined manually (Yushkevich et al. 2006).

### 2.2.3. Functional data and localisation of optic-flow selective sensory regions

In order to determine the spatial relations between the white matter tract and the optic-flow selective sensory areas (Cardin and Smith 2011; Frank et al. 2014), four regions of interest (ROIs) were identified using data acquired in separate fMRI localiser scans. The ROIs were defined as all contiguous voxels that were significantly more active with a coherent optic-flow stimulus (random dots moving along dynamically changing spiral paths) than with a random-motion stimulus (Uesaki and Ashida 2015); in the ventral intra-parietal cortex (VIP), the precuneus motion area (PcM), the junction of the intra-parietal sulcus (p2v), and the posterior region of the insular cortex (PIC+).

Functional data were acquired with a gradient echo, echo-planar sequence (39 contiguous axial slices, 3-mm isotropic voxels, TR = 2000 ms, TE = 25 ms, FoV = 192 x 192 mm^2^, flip angle = 80°, bandwidth = 2368 Hz/pixel), using a Siemens 32-channel posterior-head array coil; which gave an improved signal-to-noise ratio in the occipital cortex at the expense of the anterior part of the brain. Each subject underwent two fMRI scans. Functional data were then preprocessed and analysed with BrainVoyager QX (version 2.6, Brain Innovation, Maastricht, the Netherlands). Stimulus design and analysis methods are described elsewhere (Uesaki and Ashida 2015).

### 2.3. Data acquisition and preprocessing: Human Connectome Project dataset

Diffusion-weighted MRI data obtained from four subjects by the Human Connectome Consortium (Van Essen et al. 2013) were also analysed in this study. These data were acquired with multiple b-values (1000, 2000 and 3000 s/mm^2^) and a spatial resolution of 1.25 x 1.25 x 1.25 mm^3^. Measurements from the 2000 s/mm^2^ shell were extracted from the original dataset and used for analyses, because the current implementation of LiFE only accepts single-shell dMRI data. Human Connectome Project (HCP) data were preprocessed by the Consortium using methods that are described in Sotiropoulos et al. (2013).

### 2.4. Data analysis

#### 2.4.1. Coregistration of functional and diffusion MR images to T1-weighted images

T1-weighted 3D anatomical images were aligned to the anterior commissure-posterior commissure (AC-PC) plane. To do this, the locations of AC and PC were manually defined in the T1-weighted images. These landmarks were then used to apply rigid-body transformation to align the anatomical images to the AC-PC plane.

Preprocessed fMRI and dMRI data were coregistered to the T1-weighted images in the AC-PC space, using a two-stage coarse-to-fine approach. This process enabled a direct spatial comparison between the tract endpoints and the ROIs within the same coordinates in each subject, in the later analyses.

#### 2.4.2. Generation and optimisation of white matter connectomes

Ensemble tractography (Takemura, Caiafa, et al. 2016) was used to estimate the white matter tracts from dMRI data in each subject. Candidate streamlines were generated with mrTrix (Tournier et al. 2012), using CSD-based probabilistic tractography (step size = 0.2 mm; maximum length = 200 mm; minimum length = 10 mm; FOD amplitude stopping criterion = 0.1; vector specifying the initial direction = 20 deg). We used the entire grey matter-white matter interface region as a seed (Smith et al. 2012), and the seed voxels were randomly selected from the mask for generating candidate streamlines. Tracking was terminated when a streamline reached outside the white matter mask. We used four different angle threshold settings (angle threshold = 5.7, 11.5, 23.1, 47.2 deg). Two million candidate streamlines were generated for each curvature parameter setting. We then selected those located within the white matter posterior to the most anterior optic-flow selective area (the cingulate sulcus visual area; CSv; Cardin & Smith 2010) in each hemisphere, which were used in all subsequent analyses.

In order to obtain an optimised connectome model including streamlines generated with different curvature thresholds, we used the preselection method in ensemble tractography (Takemura, Caiafa, et al. 2016). Firstly, we separately optimised each connectome generated with a single curvature parameter using linear fascicle evaluation (LiFE; Pestilli et al. 2014; http://francopestilli.github.io/life/). We then selected 150,000 streamlines that contributed meaningfully to predicting the diffusion signals from each single-parameter connectome, and combined them to create a new candidate connectome (ensemble tractography connectome; ETC; 600,000 candidate streamlines) in each hemisphere. Finally, LiFE was applied again to optimise the ETC. As a result, the optimised ETC included 103,664 streamlines on average per hemisphere. The optimised ETC has been shown to perform better than conventional connectome models, in terms of model accuracy and anatomical representation (Takemura, Caiafa, et al. 2016; https://github.com/brain-life/ensemble_tractography). Optimised ETCs were used for identification and validation of SPIS.

#### 2.4.3. Tract segmentation

We used the cortical ROIs defined by Freesurfer segmentation (Desikan-Killiany atlas; Desikan et al. 2006) to identify the white matter tract connecting the superior and inferior parts of the parietal cortex in the optimised connectome. For the main analysis, two cortical ROIs, one in the superior parietal cortex and the other in the inferior parietal cortex were created. The ROI in the superior part of the parietal cortex was defined as a combination of two Freesurfer ROIs, “precuneus” and “superior_parietal”, because these two ROIs combined covered the positions of the functionally identified areas located in the superior parietal cortex (VIP, PcM, p2v). The Freesurfer ROI labelled “supramarginal gyrus” was used as the ROI in the inferior parietal cortex, which covered the general region including PIC+.

The tract between the superior and inferior parts of the parietal cortex was identified as a group of streamlines having one of the endpoints near the superior parietal ROI and the other near the inferior parietal ROI (within 3 mm from each grey matter ROI). The segmented tract was then refined by removing outlier streamlines. Those were streamlines that met the following criteria: (1) streamlines longer than the mean streamline length of the tract by ≥ 3 sd; (2) streamlines shorter than 15 mm; (3) streamlines of which position is ≥ 3 sd away from the mean position of the tract (Yeatman, Dougherty, Myall, et al. 2012).

The MATLAB code used to identify SPIS in this study is available at GitHub ([URL will be provided upon acceptance]).

#### 2.4.4. Estimating cortical endpoints of the tract

Streamlines terminate at the boundary between the white matter and grey matter. In order to estimate SPIS endpoints in the cortical surface representation, the coordinates of SPIS endpoints were collected, and the distance between those coordinates and the grey matter voxels was calculated. For each grey matter voxel, the number of SPIS endpoints falling within a threshold distance (3 mm) was counted. The normalised endpoint counts are plotted on the inflated cortical surface in Figure 1B. The same method was used in Takemura, Rokem et al. (2016).

**Figure 1.**
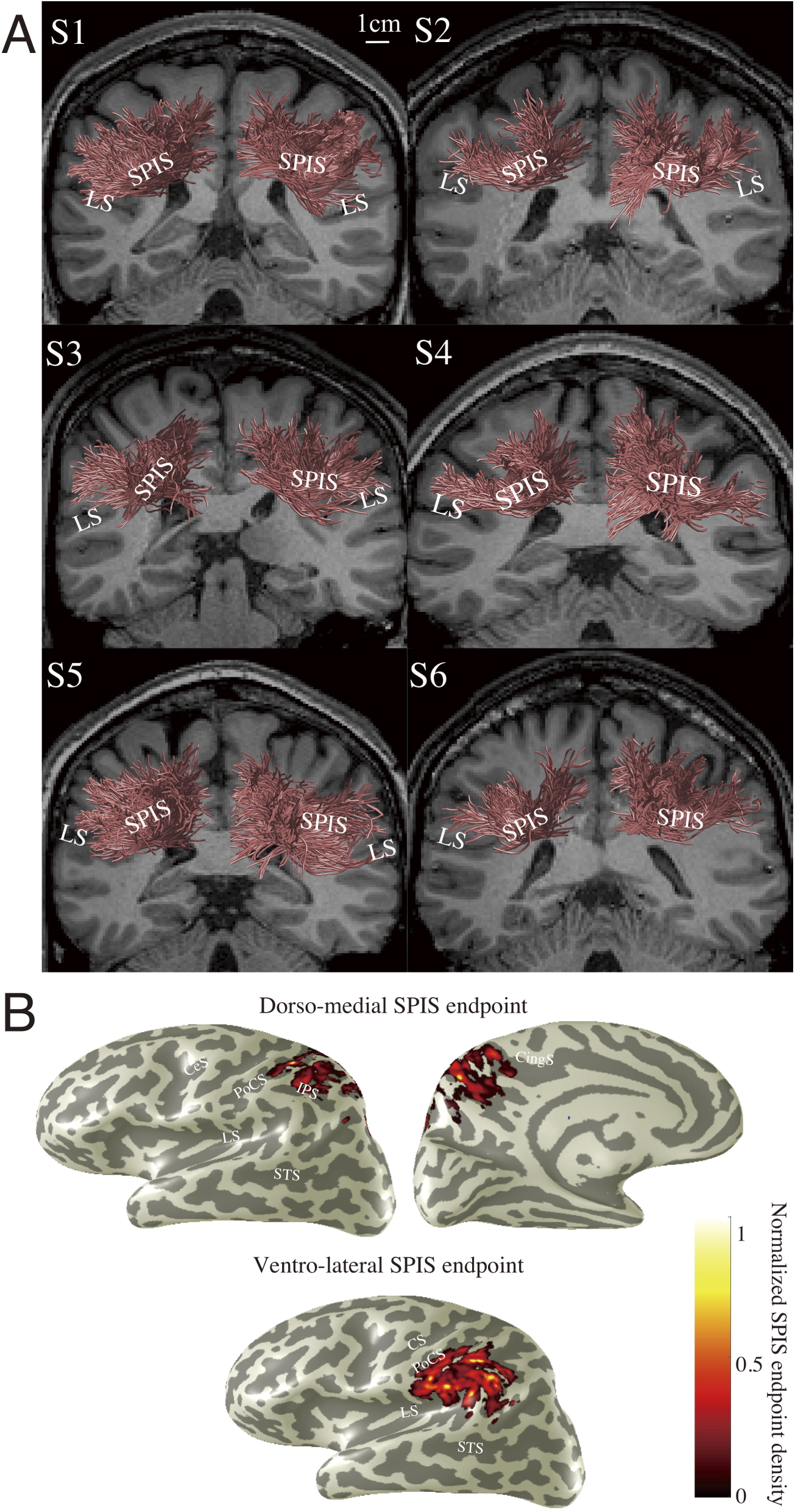
Human stratum proprium of interparietal sulcus (SPIS) estimated by tractography performed on dMRI data (KU dataset). **A.** Coronal view of SPIS (red) in 6 subjects (S1-S6; KU dataset), identified by tractography (see Materials and Methods). Scale bar (white) in the S1 panel indicates 10 mm. The background anatomical (T1-weighted) slice is located immediately anterior to the position of SPIS. LS: Lateral Sulcus. **B.** SPIS endpoints overlaid on the cortical surface (S1, left hemisphere). The spatial distance between SPIS endpoints and grey matter voxels was calculated in order to plot the number of SPIS streamlines having endpoints close to grey matter voxels (see Materials and Methods). CS: Central Sulcus, PoCS: Postcentral Sulcus, IPS: Intraparietal Sulcus, STS: Superior Temporal Sulcus, CingS: Cingulate Sulcus.

#### 2.4.5. Virtual lesion analysis

We conducted virtual lesion analysis (Pestilli et al. 2014; Leong et al. 2016; Takemura, Rokem, et al. 2016) on KU dataset to evaluate the statistical evidence supporting the existence of the white matter pathway connecting the superior and inferior parts of the parietal cortex. For this analysis, we divided the dMRI data (KU dataset) into two sessions; one was used for performing tractography, and the other was used for computing cross-validated model prediction accuracy.

Two connectome models were used in the analysis; the optimised connectome and a lesioned connectome with the streamlines of interest (the streamlines that belong to the tract connecting the superior and inferior parts of the parietal cortex) removed. We computed prediction accuracy (root mean squared error; RMSE) of those models in predicting the diffusion signals. The set of dMRI data from the second session was used as the measured diffusion signals for cross-validation. Model accuracy is described as a ratio of RMSE (*R*_*rmse*_), and it represent prediction accuracy of each model with respect to test-retest reliability (for calculation methods, see Rokem et al. 2015; Takemura, Caiafa, et al. 2016). *R*_*rmse*_ = 1 indicates that the model accuracy for predicting the diffusion signals in the second dataset equals to test-retest reliability of the diffusion signals in the same voxel.

*R*_*rmse*_ was compared in all voxels touched by the lesioned streamlines (the streamlines that belong to SPIS). The complete set of streamlines that contribute to the prediction of the diffusion measurements in those voxels is called the path-neighborhood of SPIS (Wedeen et al. 2012). This path-neighborhood includes SPIS itself and all of the other streamlines that pass through the voxels SPIS passes through. We calculated the distribution of *R*_*rmse*_ values in SPIS voxels with the entire path-neighborhood included and removed SPIS, to figure out the weight of each of the remaining streamlines.

Evidence supporting SPIS was assessed using two different measures. We compared the two *R*_*rmse*_ distributions using the strength of evidence, *S,* which is the difference in the mean *R*_*rmse*_ divided by the joint standard deviation (for technical detail, see Pestilli et al. 2014).

#### 2.4.6. Spatial proximity between SPIS endpoints and functionally-defined ROIs

In order to characterise the spatial proximity between the optic-flow selective ROIs and SPIS, we measured the proportion of grey matter voxels in each ROI (VIP, PcM, p2v and PIC+) located close to SPIS endpoints. We computed the three-dimensional distance between the endpoint of each SPIS streamline and each grey matter voxel included in the ROIs. We then calculated the proportion of voxels in each ROI located within specific distance (thresholded at 3 or 4.5 mm in volumetric space) from any SPIS streamline endpoints. This procedure is based on that reported in Takemura, Rokem, et al. (2016).

We note that there are limitations to this analysis, due to the general difficulty in associating streamline endpoints and the grey matter surface (Reveley et al. 2015), particularly for dMRI data with a standard resolution as used in this study. The aim of this analysis was to examine the general spatial proximity between the tract endpoints and functionally defined ROIs, rather than to determine the projection pattern of the tract on the cortical surface.

## 3. Results

The primary aim of this study was to identify a white matter tract connecting the superior and inferior parts of the parietal cortex. By performing tractography on the dMRI data, we successfully identified SPIS in all subjects. We compared the tractography estimates with SPIS reported in the post-mortem fibre dissection studies (Sachs 1892; Vergani et al. 2014), and evaluated the statistical evidence for the estimates (Pestilli et al. 2014) as well as consistency across datasets. The tract was identified consistently across subjects and across datasets, and seems to correspond with the tract referred to as SPIS in previous fibre dissection studies (Sachs 1892; Vergani et al. 2014). Furthermore, we assessed the spatial relations between SPIS and the optic-flow selective sensory areas localised using fMRI.

### 3.1. Anatomy of human SPIS

### 3.1.1. Tract trajectory and length

We analysed the dMRI data (KU dataset) using Ensemble Tractography (Takemura, Caiafa, et al. 2016b; see Materials and methods), whereby a whole-brain structural connectome was generated using multiple parameter settings, and then optimised using Linear Fascicle Evaluation (LiFE; Pestilli et al. 2014; see Materials and methods). We identified a white matter tract having one endpoint near the superior parietal cortex and another near the inferior parietal cortex, based on the grey matter ROIs defined by Freesurfer (Fischl 2012; see Materials and Method).

Figures 1A shows the group of streamlines that comprises the pathway we identified in each hemisphere in 6 subjects from KU dataset, superimposed on a coronal slice. The estimated tract, SPIS, primarily connects the superior and inferior parts of the parietal cortex and wraps around the intraparietal sulcus (IPS). In all subjects, SPIS in the two hemispheres are symmetrically oriented. The mean length of the estimated SPIS streamlines was 4.69 cm (SD = 0.59, N = 12 hemispheres). Figure 1B shows the estimated SPIS endpoint on the cortical surface representation in one representative hemisphere (S1, left hemisphere). Most of the dorso-medial SPIS endpoints are in the parietal areas superior to the IPS, but also in the medial areas such as the precuneus. The ventro-lateral endpoints of SPIS are in the supramarginal gyrus, parietal operculum and the posterior end of the lateral sulcus.

Additionally, we analysed the the publicly available Human Connectome Project dataset (HCP dataset; Van Essen et al. 2013) in order to assess the consistency of SPIS results across datasets. Figure 2A demonstrates SPIS in four subjects from HCP dataset, identified using the identical methods to those used to identify the tract in KU dataset. The estimated position and trajectories of SPIS are consistent with those in KU dataset (Figures 1A). Figure 2B shows a Principal Diffusion Direction (PDD) map illustrating the position of SPIS in one representative HCP subject (subject S7). PDD is often used to identify the major white matter pathways, and it allows for tract identification independent of the selection of tractography methods (Pajevic and Pierpaoli 1999; Wakana et al. 2004; Yeatman et al. 2013; Takemura, Rokem, et al. 2016). The PDD map of the HCP dataset clearly shows the existence of tracts travelling between the medial side of superior parietal cortex, and lateral inferior regions around parietal operculum and posterior part of lateral sulcus.

**Figure 2.**
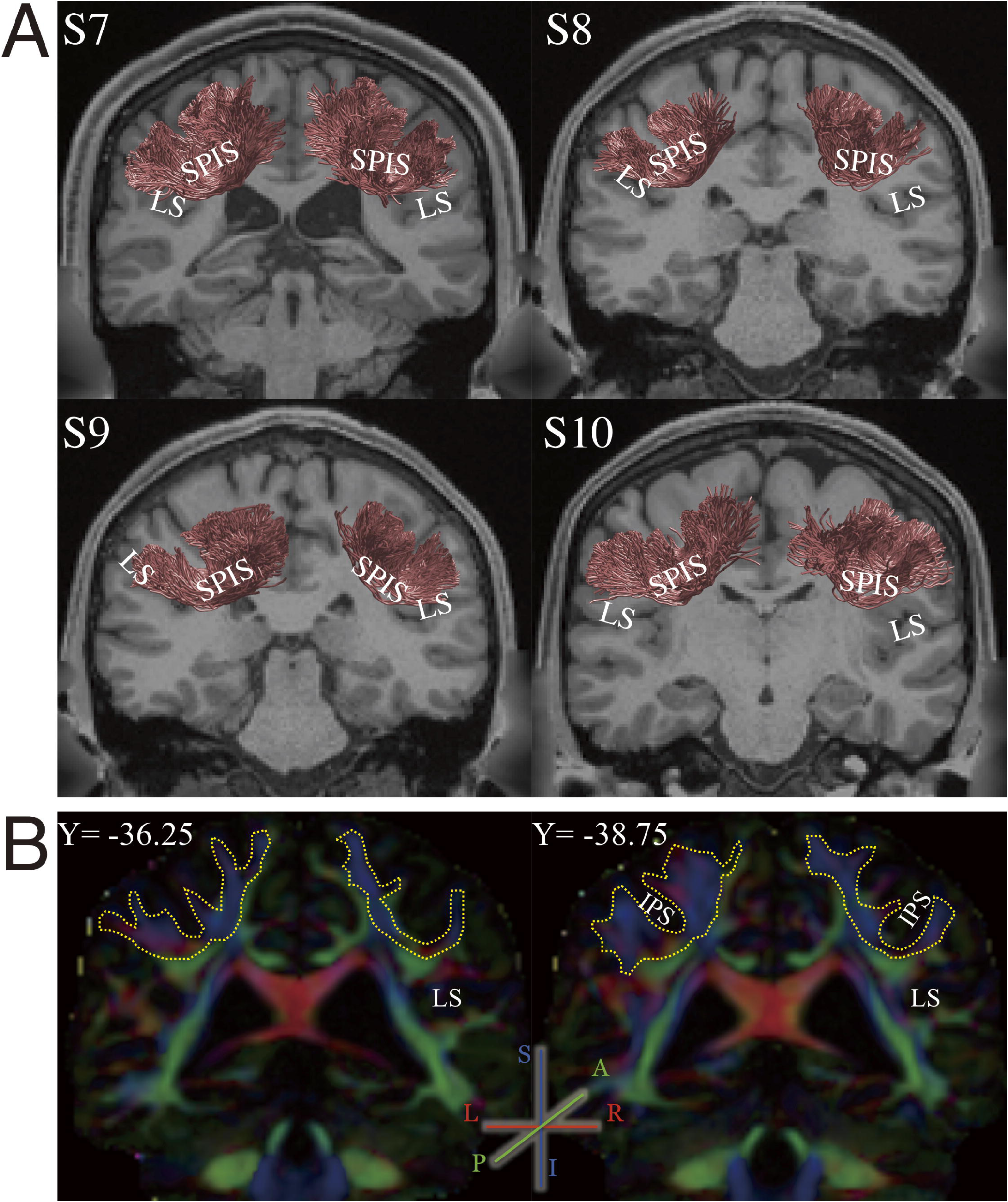
SPIS identified in HCP dataset. **A.** Coronal view of SPIS (red) in four subjects (S7-S10) from HCP dataset, identified by tractography (see Materials and Methods). The conventions are identical to those in Figure 1A. **B.** Position of SPIS highlighted in PDD map (S7, two representative coronal slices; the position of each slice is shown in ACPC coordinate). The colour scheme depicts the PDD in each voxel (blue: superior-inferior; green: anterior-posterior; red: left-right). White matter portion connecting the dorso-medial and ventro-lateral regions wrapping around the intraparietal sulcus (IPS) that is predominantly blue/purple clearly illustrates the trajectory of SPIS. Yellow dotted line highlights the position of SPIS.

The results of tractography, which is consistent across subjects and datasets, as well as voxelwise evidence of SPIS that is not based on tractography, further corroborate the evidence for SPIS.

### 3.1.2 Comparison with fibre dissection studies

We compared the anatomical position and shape of SPIS identified from dMRI data with post-mortem fibre dissection studies. SPIS has been documented in two previous post-mortem fibre dissection studies; in the classical work by a German neurologist Heinrich Sachs (1892) and more recently by Vergani and colleagues (2014). Sachs referred to this tract as *stratum proprium fissurae interparietalis* in his report, which was later rephrased by Vergani and colleagues (2014) as *stratum proprium of interparietal sulcus (SPIS)*. We adopt this term, SPIS, to refer to the tract estimated in the present study.

Figure 3 compares the position of SPIS in Sachs’s study (Figure 3A and 3B), in Vergani’s study (Figure 3C), with the pathway we identified using tractography (Figure 3D). SPIS identified in this study is consistent with SPIS reported in the fibre dissection studies in several aspect. In terms of its spatial relations with cortical landmarks, SPIS wraps around the intraparietal sulcus, and connects the parietal cortex and the dorsal bank of the lateral sulcus. The position of SPIS on the coronal slice is also consistent. The coronal slice of the anatomical image onto which SPIS estimated by tractography is superimposed in Figure 3D was chosen carefully so that it corresponded with the slices used in the fibre dissection studies as closely as possible. Whilst it is not possible to perfectly match the position of the slice between our MRI data and fibre dissection studies, it is qualitatively consistent across the four presentations in Figure 3, in terms of the positions of the sulci (i.e. the lateral sulcus and the intraparietal sulcus) and the lateral ventricle. Although SPIS in the three studies cannot be compared quantitatively due to the difference in the methodology used, Figure 3 demonstrates that the position and trajectory of SPIS identified with our dMRI data agrees with those of SPIS reported in the fibre dissection studies.

**Figure 3.**
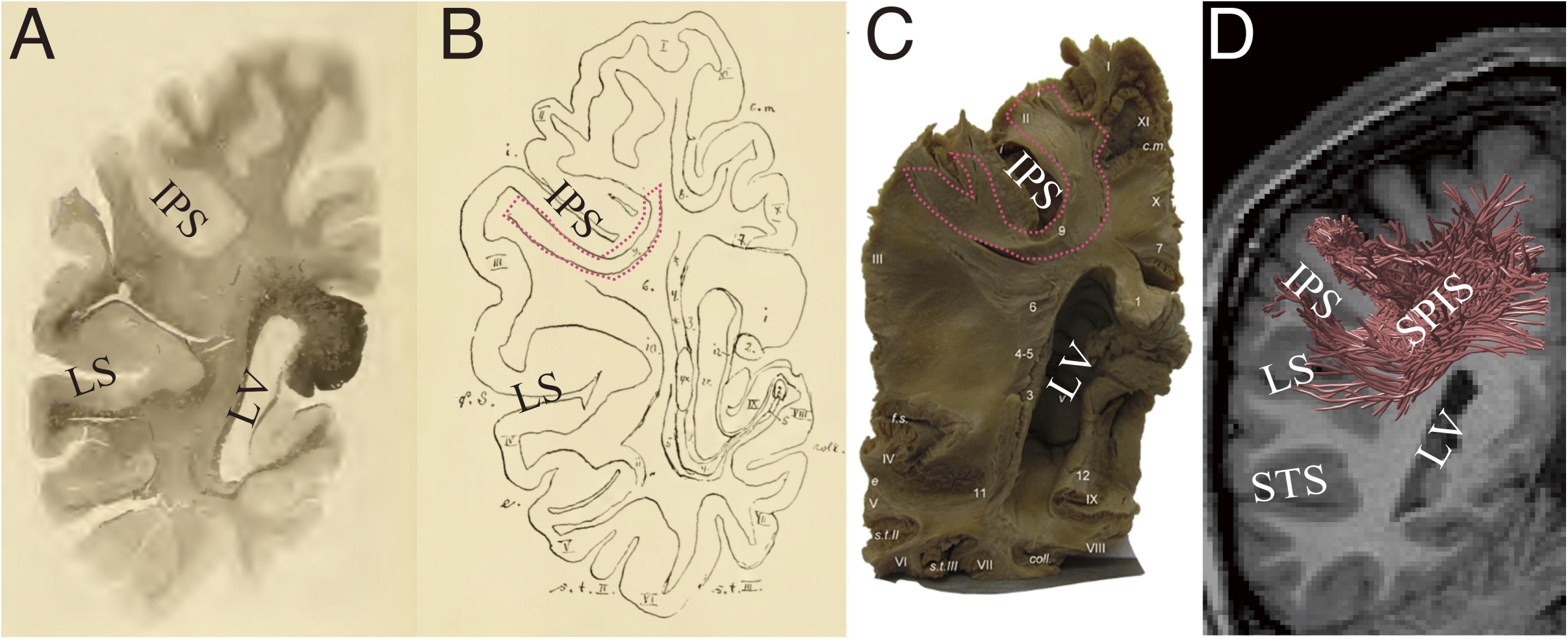
Tractography results are consistent with classical and modern fibre dissection studies. **A.** White matter tract referred to as SPIS on a coronal histological slice and **B.** SPIS (red outline) in the schematic diagram of the fibres visualised in the photo in Panel A (Sachs 1892). Sachs (1892) noted that this slice is approximately 75 mm anterior to the occipital pole (Forkel et al. 2015). **C.** SPIS identified in a post-mortem human brain in the modern fibre dissection study (right hemisphere; Vergani et al. 2014). The position of SPIS is highlighted with red outline. Reproduced from Vergani et al. (2014) with permission. **D.** SPIS estimated by tractography (red lines) in one representative hemisphere (S1; right hemisphere). The image has been flipped (for the original image, see Figure 1A) so that the slice corresponds with the fibre dissection studies. The background coronal slice (ACPC coordinate, Y = -44; approximately 65 mm anterior to the occipital pole) is located immediately anterior to the estimated SPIS. The position and the trajectory of SPIS are qualitatively consistent with those of SPIS reported in the fibre dissection studies (Panels A-C). LV: Lateral Ventricle, STS: Superior Temporal Suclus, IPS: Intraparietal Sulcus.

### 3.1.3. Position of SPIS with respect to major white matter tracts

Figure 4 shows SPIS overlaid on a sagittal plane of the T1-weighted image, along with other major white matter tracts reported in previous studies; the arcuate fasciculus (AF; Wakana et al. 2004), posterior arcuate (pArc; Catani et al. 2005; Weiner et al. 2016) and vertical occipital fasciculus (Yeatman et al. 2013; Yeatman, Weiner, et al. 2014; Duan et al. 2015; Takemura, Rokem, et al. 2016) in one representative subject (S1).

**Figure 4.**
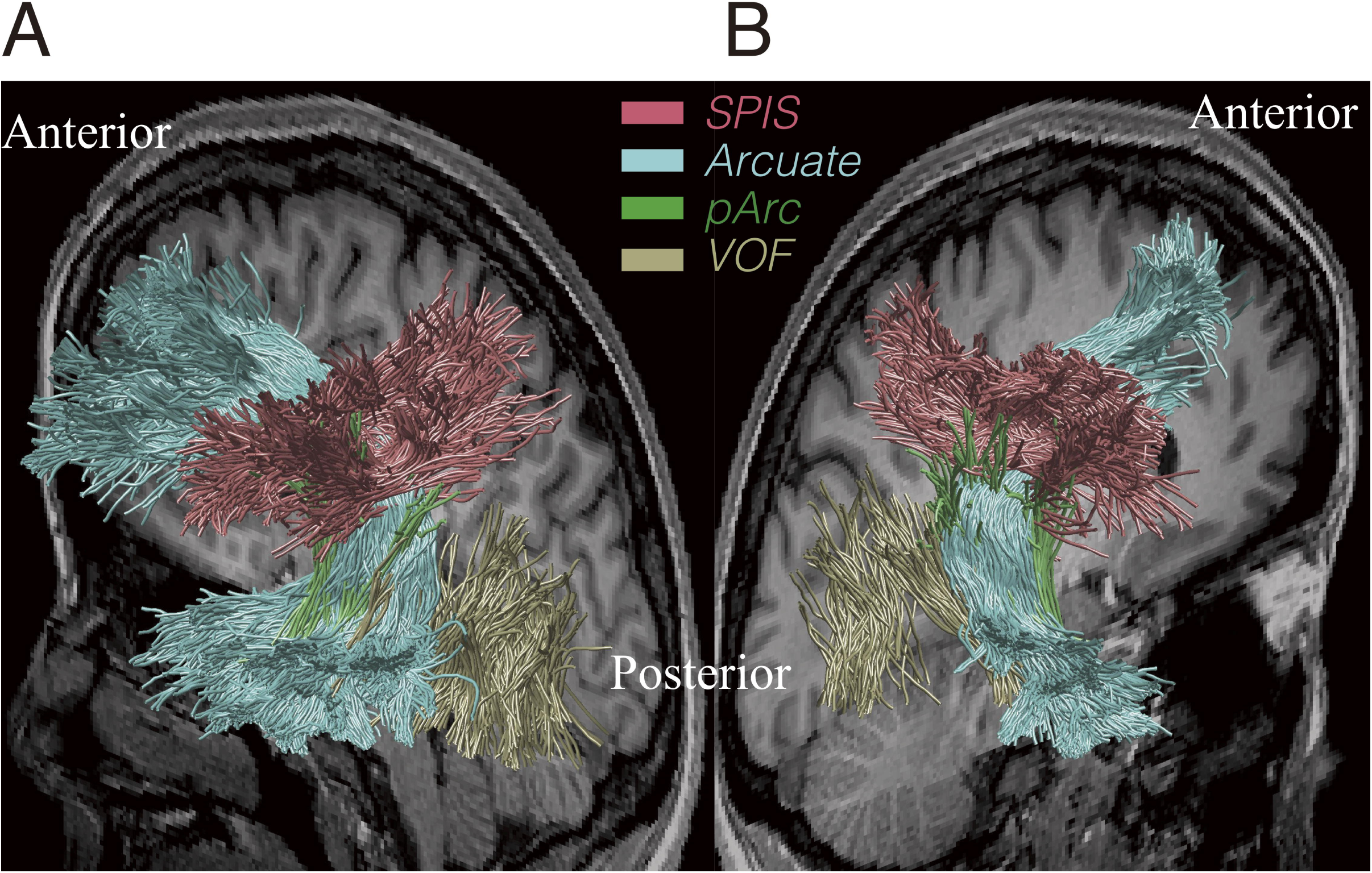
Position of SPIS with respect to other tracts. **A.** Position of SPIS with respect to other tracts in the left hemisphere of one representative subject (S1). The background T1-weighted image is a sagittal slice in the medial portion of the brain. SPIS (red) is located superior to the vertical occipital fasciculus (VOF; yellow) and posterior arcuate (pArc; green). SPIS lies on the superior surface of, and crosses with the arcuate fasciculus (Arcuate; light blue). **B.** Position of SPIS with respect to other tracts in the right hemisphere in the same subject (S1).

SPIS is located adjacent to AF; in fact, SPIS intersects with the dorsal surface of AF. This crossing may be one of the reasons that this tract has been relatively neglected in the literature, as resolving crossing fibres is one of the critical limitations of the diffusion tensor-based approach (Frank 2001; Tournier et al. 2012). Interestingly, the intersection between SPIS and AF may explain the pattern of previous dMRI results along AF. Yeatman and colleagues (2011) investigated the fractional anisotropy (FA; Basser and Pierpaoli 1996) along AF, and found that there is a large dip in the FA value along the length of the fasciculus in the vicinity of the temporal cortex. Yeatman et al. (2011) suggests that this dip is partially accounted for by the sharp curvature of AF, but also by partial voluming with crossing fibres. Since the location of this dip along the trajectory of AF coincides with the position of SPIS, it seems plausible that this is where the AF intersects with SPIS.

SPIS is also located near pArc, but the trajectory and endpoints of SPIS are distinct from pArc, which connects the parietal cortex and the anterior inferior temporal cortex.

Although there are a few neighbouring tracts, some of which cross/kiss SPIS, the differences in the trajectory and locations of endpoints between those known tracts and SPIS clarify that SPIS is a distinct tract.

### 3.2. Statistical evidence in support of SPIS

To evaluate the strength of statistical evidence supporting the existence of SPIS, we used virtual lesion methods (Honey and Sporns 2008; Pestilli et al. 2014; Leong et al. 2016; Takemura, Rokem, et al. 2016). We first computed the cross-validated prediction accuracy for diffusion signal (Ratio of RMSE; *R*_*rmse*_; Rokem et al. 2015; Takemura, Caiafa, et al. 2016) in models with lesioned and unlesioned SPIS. We then compared the distribution of *R*_*rmse*_ of the two models to predict the diffusion signals within SPIS voxels (see Materials and Methods for details). We quantify the strength of evidence (*S*) in support of the SPIS by calculating the difference of *R*_*rmse*_ in lesioned and unlesioned model divided by the standard deviation of the *R*_*rmse*_ (Pestilli et al., 2014).

Figure 5A describes the mean and variance of the statistical evidence for SPIS across subjects, yielded by virtual lesion analysis. The mean strength of statistical evidence for SPIS was *S* = 76.25 (*SD* = 10.87) for the left hemisphere, and S = 84.7 (SD = 6.72) for the right hemisphere. Figure 5B describes the two-dimensional histogram of *R*_*rmse*_ in the SPIS lesioned and unlesioned models for the left hemisphere in one representative subject (S1). In many voxels, the SPIS lesioned model showed substantially lower model accuracy (higher *R*_*rmse*_) as compared with the unlesioned model, indicating that SPIS is necessary to explain the diffusion signals within those voxels. Thus, in addition to the results of tractography and their consistency with the findings of previous post-mortem studies at visual inspection, there is strong statistical evidence supporting the existence of SPIS.

**Figure 5.**
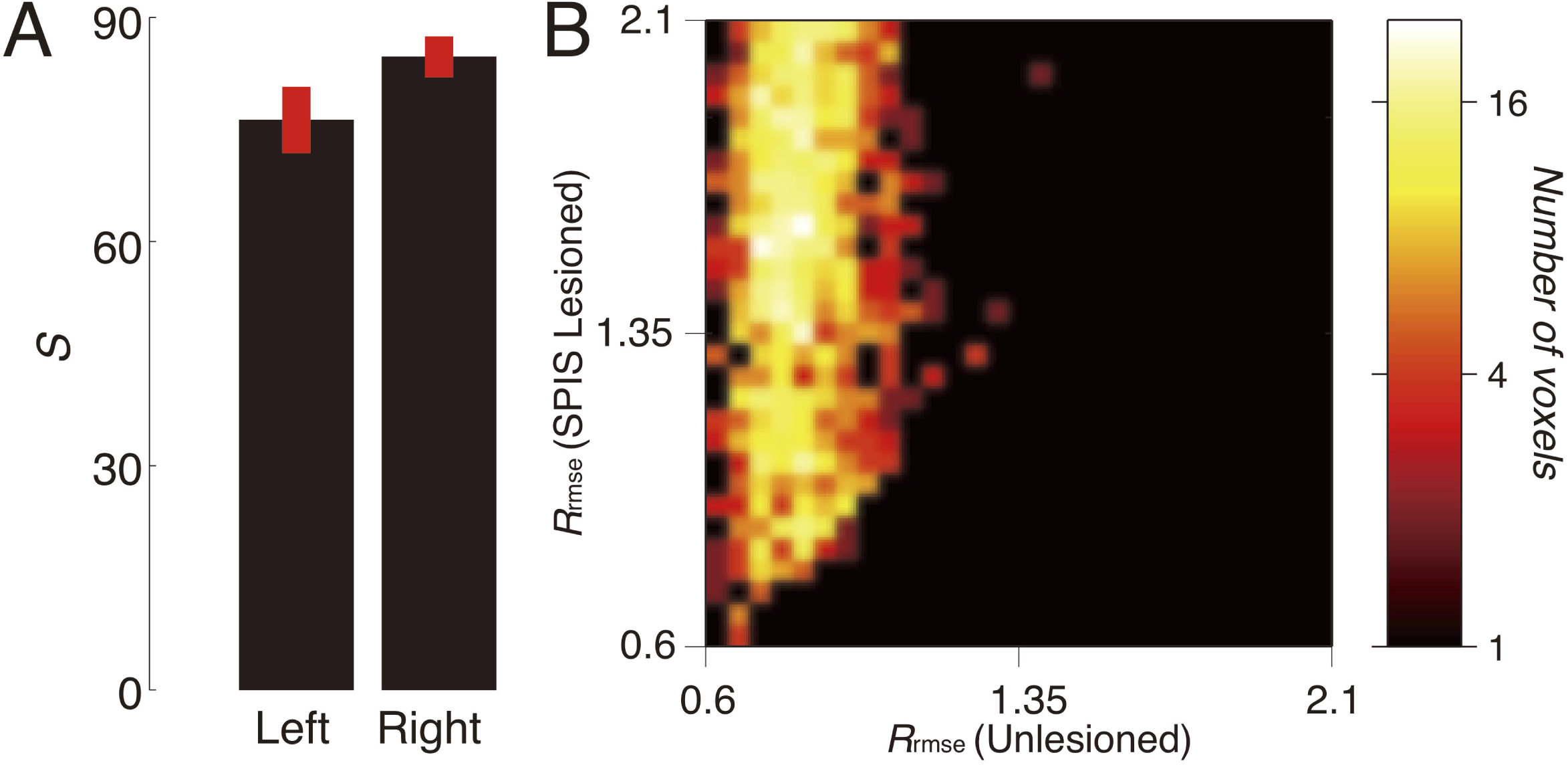
Statistical evidence in support of SPIS. **A.** Mean *S* in support of left and right SPIS across subjects. Error bars depict ±1 s.e.m across subjects. **B.** Two-dimensional histogram comparing the model accuracy (*Rrmse*) between the lesioned and unlesioned models (horizontal axis: unlesioned model; vertical axis: lesioned model) for SPIS in the left hemisphere in one representative subject (S1). Prediction accuracy is substantially lower with the lesioned model. Colour bar (right panel) indicates the number of voxels.

### 3.3. SPIS and its relations with optic-flow selective cortical areas

With the subjects in KU dataset, we further conducted fMRI experiments to localise cortical sensory areas selective for optic-flow stimulation in order to examine the spatial proximity of SPIS endpoints and those functionally-defined areas.

### 3.3.1. Functional localisation

To localise the cortical areas selective for optic-flow stimulus, blood-oxygen level dependent (BOLD) responses to the coherent optic-flow stimulus was contrasted against those to the random-motion stimulus. We identified four of the cortical areas known to be selective for optic flow (Figure 6; VIP, p2v, PcM, PIC+; Cardin and Smith 2011; Uesaki and Ashida 2015). Areas VIP, p2V and PcM are located in the superior part of the parietal cortex, and PIC+ in the posterior end of the lateral sulcus. The locations of those areas in Talairach coordinates were consistent with those of the corresponding areas reported in previous studies (Cardin and Smith 2011; Frank et al. 2014; Uesaki and Ashida 2015). All four areas were successfully identified in nine hemispheres. In other three hemispheres, either PcM or VIP was not identified.

**Figure 6.**
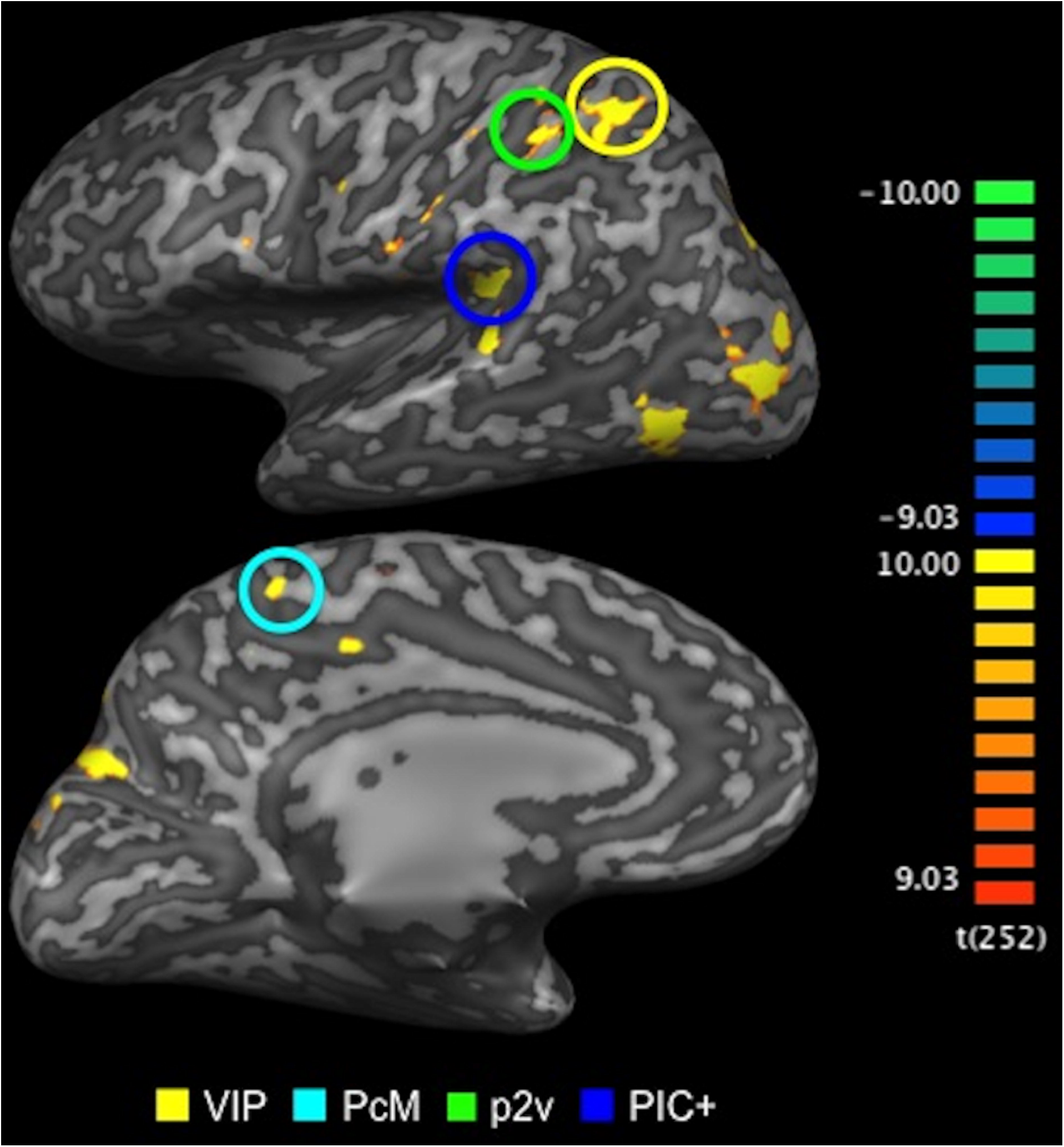
Optic-flow selective areas localised using fMRI. Cortical areas that showed significantly greater BOLD responses to optic-flow stimulus than to random-motion stimulus (*p* < .005, uncorrected). Activation maps are superimposed on the inflated cortical surface of the left hemisphere in one representative subject (S1). Colour-coded bar (right panel) indicates statistical t-values. Four of the cortical areas selective for optic flow (VIP, PcM, p2v and PIC+) were successfully identified.

### 3.3.2. SPIS endpoints and optic-flow selective cortical areas

Subsequently, we examined the spatial proximity between the cortical areas selective for optic flow (Figure 6) and SPIS endpoints. Although there is a limitation to use tractography for identifying the tract endpoint in grey matter (Reveley et al. 2015), it is still useful to understand how closely functionally-defined ROIs are located to the tract endpoints in order to infer any potential implication of the tract in information transmission during optic-flow processing. We analysed the general spatial proximity between SPIS endpoints and the optic-flow selective areas (VIP, PcM, p2v and PIC+).

Figure 7A depicts the relative position of SPIS with respect to the cortical areas selective for optic flow, in the left hemisphere of one representative hemisphere subject (S1, left hemisphere). Three of the optic-flow selective areas (VIP, p2v and PcM) in the parietal cortex are located near the dorso-medial endpoint of SPIS. PIC+ is located in the posterior end of the lateral sulcus, near the ventro-lateral endpoint of SPIS. Figure 7B summarises the proportion of grey matter voxels in each optic-flow selective area located near SPIS endpoints (see Materials and Methods). Approximately 40 and 80% of voxels in each grey matter ROI are located in the vicinity of SPIS endpoints, depending on the distance threshold for defining the spatial proximity between grey matter voxels and tract endpoints (i.e. proximity was thresholded at 3 mm or 4.5 mm). It seems highly likely that SPIS is part of the anatomical connection between the optic-flow selective areas in the superior parietal cortex (VIP, p2v and PcM) and the PIC+.

**Figure 7.**
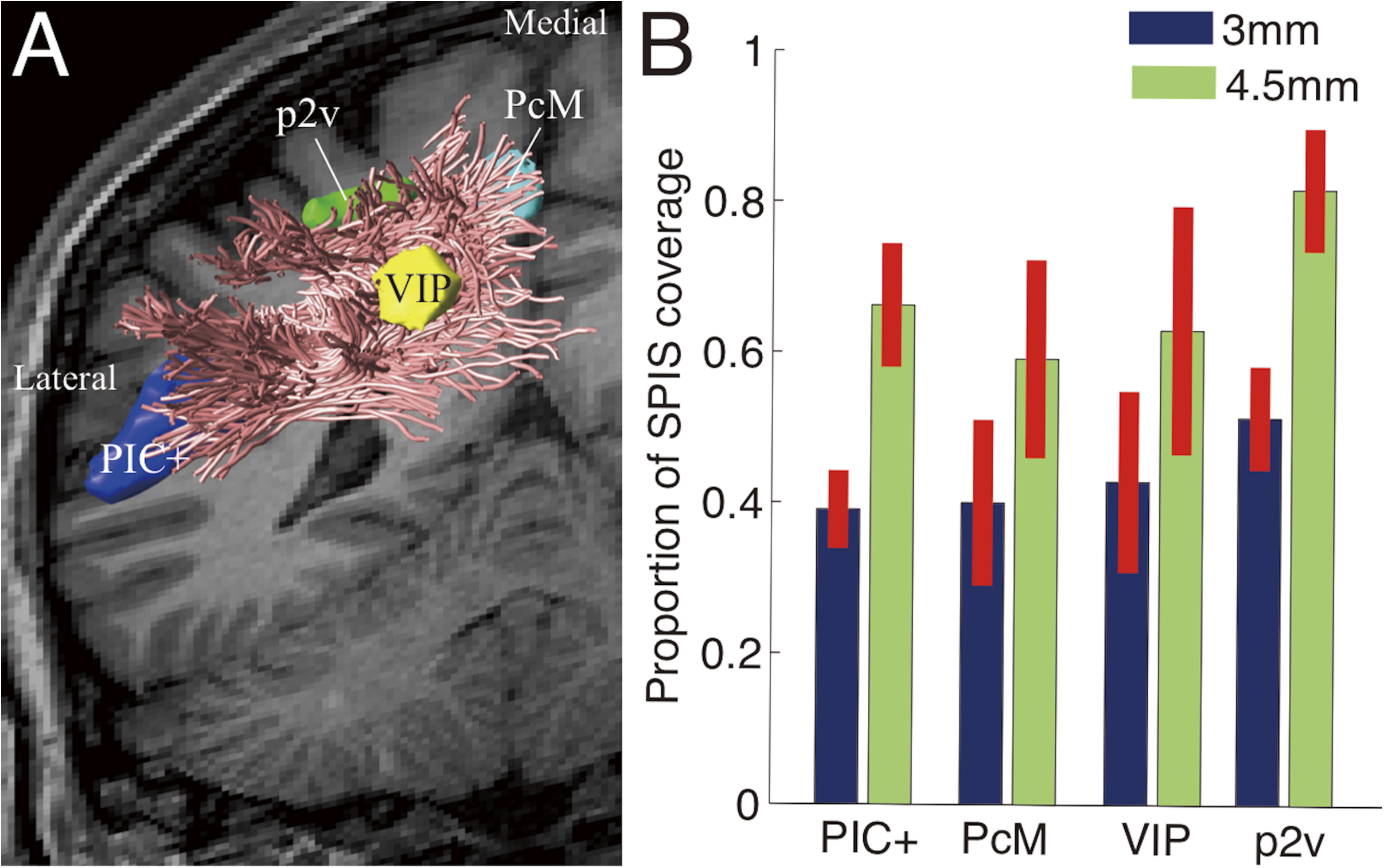
Spatial proximity between SPIS endpoints and cortical ROIs selective for optic-flow stimulation. **A.** Position of SPIS (magenta lines) in relation to the cortical ROIs identified by optic-flow stimulation using fMRI (dark blue: PIC+; light blue: PcM; yellow: VIP; green: p2v) in the left hemisphere in one representative subject (S1). **B.** SPIS map coverage across all hemispheres. Vertical axis represents the proportion of voxels in each ROI within 3 mm (blue) and 4.5 mm (green) from SPIS endpoints. Error bars indicate ± 1 s.e.m. across hemispheres.

## 4. Discussion

*Stratum proprium of interparietal sulcus* (SPIS) was originally discovered in a post-mortem fibre dissection study by Sachs (1892) and was reproduced in another post-mortem study by Vergani et al. (2014). Here, SPIS was successfully identified in the living human brain for the first time, using dMRI and tractography.

### 4.1. Comparison of SPIS results with anatomical studies

In this study, we investigated a white matter tract that has been largely overlooked in the visual and cognitive neuroscience, SPIS, using dMRI-based tractography and fascicle evaluation techniques. In spite of the challenges of using tractography to study a novel white matter tract, SPIS was consistently identified across subjects and datasets. Between our dMRI results and the findings of the following anatomical studies, there is converging evidence supporting the existence of SPIS.

### 4.1.1. Human post-mortem fibre dissection studies

Most importantly, our results are consistent with human post-mortem fibre dissection studies (Figure 3; Sachs 1892; Vergani et al. 2014). In those studies, SPIS was found to be located immediately posterior to the central sulcus, wrapping around the intraparietal sulcus, and to range between the superior parietal cortex and the lateral fissure. The position and the trajectory of SPIS reported in the post-mortem fibre dissection studies are consistent with those of SPIS identified *in vivo* in two independent dMRI datasets in this study (Figures 1 and 7).

To our knowledge, the first description of SPIS appeared in the atlas of Heinrich Sachs (1892); a German neurologist and neuroanatomist who studied under Wernicke. Sachs’s atlas (1892) describes the white matter tracts in the human post-mortem brain in great detail, including the U-fibre system which has not been studied extensively in the living human brain. One of the short-association tracts described is a tract termed *stratum proprium fissurae interparietalis*. Despite its relevance to perceptual neuroscience, Sachs’s atlas has been largely overlooked in the literature partly due to the lack of translation of the atlas from German to English (see Forkel et al. 2015; for a historical review and English translation of the atlas). SPIS documented in this classical atlas was reproduced in a modern fibre dissection study by Vergani and colleagues (2014).

Our results describe the characteristics of SPIS identified in the living human brain, using modern neuroimaging techniques, which are highly consistent with the findings of the human post-mortem studies; hence providing further evidence for SPIS.

### 4.1.2. Macaque tracer study

Additionally, we note that a tract similar to human SPIS in the macaque brain has been reported in a tracer study. In their extensive study, Schmahmann and Pandya (2006) injected retrograde tracers into the macaque brain, and inspected the trajectory of white matter tracts from the injection sites. They reported several major white matter tracts seemingly homologous to human major white matter tracts identified in dissection studies (such as the inferior longitudinal fasciculus, and the superior longitudinal fasciculus); and those findings were later substantiated by macaque dMRI results (Schmahmann et al. 2007). In addition to the major white matter tracts, Schmahmann and Pandya (2006) also reported a fibre bundle wrapping around the intraparietal sulcus. They note (page 120):

> *A dorsal fiber bundle lies subjacent to the cortex of the lower bank of the IPS and terminates in a columnar manner in area POa and in area IPd (Scs. 105, 113). These fibers continue medially and then curve around the depth of the IPS to ascend in the white matter of the superior parietal lobule. They terminate in area I in a columnar manner and then first layers of area 3b and 3a in the caudal bank and depth of the central sulcus (Sc. 105). Further caudally, these medially directed fibers terminate in a columnar manner in area 2 (Sc. 113).*

Because of the compelling similarity between this fibre bundle identified in the macaque brain and human SPIS in terms of their anatomical positions and shapes, it could be hypothesised that this fibre bundle in macaque may be the homologue of human SPIS identified in this study.

Whilst the white matter structure of the macaque brain may be different from that of the human brain to some extent (Rilling et al. 2008), the fact that there is a white matter tract in the macaque brain that largely resembles human SPIS is encouraging for future investigations on human-macaque homology with respect to SPIS. There is a growing trend in neuroanatomy to use dMRI methods to compare the macro-scale white matter anatomy of the human brain and that of the macaque brain (Schmahmann et al. 2007; Thiebaut de Schotten et al. 2011; Jbabdi et al. 2013; Mars et al. 2015; Takemura et al. in press), which complements studies that investigate human-macaque homology of cortical maps using fMRI (Tsao et al. 2003, 2008; Wade et al. 2008; Goda et al. 2014; Kolster et al. 2014). It will be beneficial to study the precise anatomy of SPIS both in humans and macaques, in order to integrate the insights from macaque electrophysiology as well as tracer studies, and human fMRI studies investigating the neuronal network for multisensory integration guiding self-motion perception.

### 4.2. Functional localisation of optic-flow selective sensory regions

Recent fMRI studies have shown that the sensory areas in the superior parietal regions (VIP, PcM, p2v) as well as an area around the posterior end of the lateral sulcus and parietal operculum (PIC+) are activated by optic-flow stimulation (Wall and Smith, 2008; Cardin and Smith 2011; Greenlee et al. 2016). As in Uesaki and Ashida (2015), this study employed the functional localiser based on that described in Pitzalis et al. (2010), in order to identify VIP, PcM, p2v and PIC+. The locations of those regions are consistent with those of the counterparts reported in previous studies (Cardin and Smith 2011; Uesaki and Ashida, 2015).

We note that the definition and terminology of the area referred as PIC+ in this study have been debated in the literature. In some earlier publications (Wall and Smith 2008; Cardin and Smith 2010, 2011; Uesaki and Ashida 2015), an area identified using optic-flow localisers was referred to as the parieto-insular vestibular cortex (PIVC) and was considered to be involved in integrating visual and vestibular information to guide self-motion perception. However, a recent vestibular fMRI study showed that PIVC is selectively responsive to vestibular stimulation, and is unlikely to be activated by visual stimulation (Frank et al. 2016; Greenlee et al. 2016). Frank and colleagues (2014) also suggested that PIVC and the area activated by visual stimulation, which is referred to as “PIC” in their study, are two independent areas. Their findings show that PIVC is purely vestibular, whilst PIC is predominantly visual but also processes vestibular information. Here, we use “PIC+” to refer to the area around the posterior end of the lateral sulcus and parietal operculum, activated during optic-flow stimulation, as we did not examine the responsiveness of the area to vestibular stimuli.

Results suggest that the ventro-lateral endpoint of SPIS is near PIC+ (Figure 7), but it is unclear whether this endpoint is also near PIVC identified in vestibular fMRI studies (Frank and Greenlee 2014; Frank et al. 2014; Greenlee et al. 2016). Considering the proximity between PIC and PIVC (Frank and Greenlee 2014; Frank et al. 2014; Greenlee et al. 2016), it is possible that the ventro-lateral endpoint of SPIS is also adjacent to PIVC. Future studies should assess whether PIVC is directly connected to the superior part of the parietal cortex through SPIS, or indirectly connected via short-range connections with PIC+, in order to construct a more comprehensive model to understand how visual and vestibular signals are transmitted between these areas to guide self-motion perception.

### 4.3. SPIS and its implication in multisensory integration

Optic flow is a moving pattern on the retina caused by the relative motion between the observer and the scene, and is one of the important visual cues to the estimation of self-motion (Gibson 1950, 1954; Warren and Hannon 1988). However, in most cases, perception of self-motion depends on integration of optic-flow information and signals from other sensory modalities like the vestibular system. In order to understand the neuronal mechanism involved in the estimation of self-motion, it is important to elucidate how the visual and vestibular signals are integrated when we observe optic flow. Previous fMRI studies investigating the cortical areas selective for optic-flow and vestibular stimuli suggest that the sensory areas in the parietal cortex are involved in visuo-vestibular integration necessary for self-motion estimation (Wall and Smith 2008; Cardin and Smith 2011; Greenlee et al. 2016). Yet, the white matter anatomy that supports the communication amongst those areas has received very little attention in the literature of visual and cognitive neuroscience, even though the existence of SPIS has been known for over a century (Sachs 1892; Vergani et al. 2014).

One of the biggest advantages of the dMRI approach is that the positions of estimated white matter tracts and functionally localised cortical areas can be compared in the brain of the same individual. This is particularly important in order to hypothesise the types of information that are transferred via the tracts of interest (Kim et al. 2006; Greenberg et al. 2012; Yeatman et al. 2013; Takemura, Rokem, et al. 2016). We combined dMRI and fMRI, and analysed the spatial proximity between the SPIS endpoints and the optic-flow selective cortical areas localised in the same subjects. Results show that the dorso-medial SPIS endpoints are near VIP, PcM and p2v, and ventro-lateral SPIS endpoints near PIC+. These cortical areas have been associated with convergence of visual and vestibular information regarding self-motion (Fetsch et al. 2009; Prsa et al. 2012; Uesaki and Ashida 2015; Kleinschmidt et al. 2002; Kovacs et al. 2008; Wiest et al. 2004; Butler et al. 2010). Our results and those findings together suggest that communication between VIP, PcM, p2v and PIC+ likely plays a crucial role in multisensory integration necessary for accurate perception of self-motion, and that it is supported by SPIS. The spatial relationship between SPIS and the optic-flow selective areas will have implications for interpreting the consequence of white matter lesions that include SPIS, or exploring the neuronal basis of individual difference in self-motion perception.

### 4.4. Future directions and limitations

Our findings show that SPIS is an important structure supporting communication amongst sensory areas in the parietal cortex. Although SPIS is discussed mainly within the contexts of multisensory integration and optic-flow processing in this article, SPIS endpoints appear to be near the cortical areas involved in other cognitive functions such as attention (Corbetta and Shulman 2002), memory (Uncapher and Wagner 2009) and body-ownership (Blanke 2012). Future dMRI studies should examine the properties of SPIS in relation to those cognitive functions, as well as development and diseases. In order to facilitate further research on SPIS, we describe the methods and provide open-source implementations ([URL will be provided upon acceptance]).

It must also be noted that it is possible that our results represent only a subset of SPIS (Oishi et al. 2008), as we used standard-resolution dMRI data (2 mm isotropic for KU dataset; 1.25 mm for HCP dataset) to estimate the trajectory of the tract. LiFE analysis yields statistical evidence on tracts supported by dMRI data, and generally supports larger portion of fibre pathways as the data resolution improves (Pestilli et al. 2014; Takemura et al. in press). Tractography based on data with higher resolutions would likely lead to extraction of a larger portion of SPIS. Likewise, estimation of cortical endpoints would be more accurate with better-quality data, as some cortical endpoints are still missed even with the best dMRI datasets currently attainable (Reveley et al. 2015).

## 5. Conclusion

This study identified a white matter tract, SPIS, in the living human brain using dMRI and tractography. It is located immediately posterior to the central sulcus and around the intraparietal sulcus; and connects the superior and inferior parts of the parietal cortex. The location and the trajectory of SPIS are consistent with those observed in post-mortem fibre dissection studies by Sachs (1892) and Vergani et al. (2014). SPIS was identified consistently across two independent dMRI datasets, and the existence of SPIS was further corroborated by statistical evidence. These findings place SPIS in a good position to channel neuronal communication between the distant cortical areas underlying visuo-vestibular integration necessary for optic-flow processing and perception of self-motion. *In vivo* identification and characterisation of SPIS using dMRI data and tractography will open new avenues to further studying this tract in relation to diseases, development and brain functions.

## Acknowledgement

This study was conducted using the MRI scanner and related facilities of Kokoro Research Center, Kyoto University; and was supported by Japan Society for the Promotion of Science (JSPS) KAKENHI (B26285165), Grant-in-Aid for JSPS Fellows (to HT and MU) and JSPS Postdoctoral Fellowship for Research Abroad (HT). HT is a JSPS Superlative Postdoctoral Fellow. We thank Franco Pestilli for comments on the earlier version of this manuscript, Brian Wandell, Lee Michael Perry, Shumpei Ogawa and Ariel Rokem for their technical advice on this study, and Michele Furlan for rendering analysis tools. We also thank Kaoru Amano, Atsushi Wada and Noburu Nushi for providing the computing environment to run the analyses. We also acknowledge the stimulated discussion in the meetings of the Cooperative Research Project of the Research Institute of Electrical Communication, Tohoku University. Data were provided in part by the Human Connectome Project, WU-Minn Consortium (Van Essen, D. and Ugurbil, K., 1U54MH091657).

